# *ERBB2*/*HOXB13* co-amplification with interstitial deletion of *BRCA1* defines a unique subset of breast cancers

**DOI:** 10.1101/2024.08.04.605361

**Authors:** Irene Rin Mitsiades, Maristela Onozato, A. John Iafrate, Daniel Hicks, Dennis C. Sgroi, Esther Rheinbay

## Abstract

**Background:** The *HOXB13/IL17BR* gene expression biomarker has been shown to predict response to adjuvant and extended endocrine therapy in patients with early-stage ER+ HER2- breast tumors. *HOXB13* gene expression is the primary determinant driving the prognostic and endocrine treatment-predictive performance of the biomarker. Currently, there is limited data on *HOXB13* expression in HER2+ and ER- breast cancers. Herein, we studied the expression of *HOXB13* in large cohorts of HER2+ and ER- breast cancers.

**Methods:** We investigated gene expression, genomic copy number, mutational signatures, and clinical outcome data in the TGGA and METABRIC breast cancer cohorts. Genomic-based gene amplification data was validated with tri-colored fluorescence *in situ* hybridization.

**Results:** In the TCGA breast cancer cohort, *HOXB13* gene expression was significantly higher in HER2+ versus HER2- breast cancers, and its expression was also significantly higher in the ER- versus ER+ breast cancers. *HOXB13* is frequently co-gained or co-amplified with *ERBB2*. Joint copy gains of *HOXB13* and *ERBB2* occurred with low-level co-gains or high-level co-amplifications (co-amp), the latter of which is associated with an interstitial deletion that includes the tumor suppressor *BRCA1*. *ERBB2/HOXB13* co-amp tumors with interstitial *BRCA1* loss exhibit a mutational signature associated with APOBEC deaminase activity, and copy number signatures associated with chromothripsis and genomic instability. Among *ERBB2*-amplified tumors of different tissue origins, *ERBB2/HOXB13* co-amp with a *BRCA1* loss appeared to be unique to breast cancer. Lastly, patients with *ERBB2/HOXB13* co-amplified and *BRCA1* lost tumors displayed a significantly shorter progression-free survival (PFS) than those with *ERBB2*-only amplifications. The difference in PFS was restricted to the ER- subset patients and this difference in PFS was not solely driven by *HOXB13* gene expression.

**Conclusions:** *HOXB13* is frequently co-gained with *ERBB2* at both low-copy number level or as complex high-level amplification with relative *BRCA1* loss. *ERBB2/HOXB13* amplified, *BRCA1*-lost tumors are strongly enriched in breast cancer, and patients with such breast tumors experience a shortened PFS.

## Background

The Breast Cancer Index (BCI) is a gene expression–based signature comprised of two functional biomarker panels, the Molecular Grade Index (MGI) and the two-gene ratio *HOXB13/IL17BR*. In patients with hormone receptor (HR) positive tumors, the *HOXB13/IL17BR* gene expression ratio has been shown to be both a prognostic and predictive biomarker for women with ER+ breast cancer^1–5^. As a prognostic biomarker, the *HOXB13/IL17RB* gene expression ratio has been shown to predict early (0-5 years, yrs), late (5-10 yrs) and overall (0-10 yrs) distant disease recurrence^4,5^, and as a predictive biomarker it has been shown to predict adjuvant and extended adjuvant endocrine therapy response across a variety of treatment scenarios^1–3,6^. *HOXB13* is the primary determinant of endocrine benefit and response^7^. In preclinical models of estrogen receptor positive (ER+) breast cancer, expression of *HOXB13* has been shown to be modulated by estradiol and tamoxifen^8^, and a recent ChIP-seq-based study has demonstrated that HOXB13 reprograms the binding pattern of the pioneer factor FOXA1 and the ER cistrome^7^. The clinical and preclinical data strongly suggest an important role of HOXB13 in ER biology.

HOXB13 is a transcription factor that belongs to the homeobox (*HOX*) gene family, an essential group of developmental transcriptional regulators that are critical for embryonic development^9–11^. In humans, there are 39 HOX genes that are divided into four different HOX clusters (A, B, C, and D) located on chromosomes 7p, 17q, 12q, and 2q, respectively^12^. The clustered organization of HOX genes is highly conserved from *Drosophila* to man and each gene within a cluster displays a pattern of expression during embryogenesis that is contingent on its relative position within its cluster^13^. Like other *HOX* gene family members, *HOXB13* expression is generally restricted to undifferentiated and/or proliferating cells during embryogenesis^11^. However, dysregulated expression of *HOXB13* has been described in endocrine-responsive tumors that include prostate, ovarian, endometrial and breast carcinomas^14–17^.

Studies investigating the expression of *HOXB13* in HER2+ and ER- breast cancer are limited. Previously, in a small single-institution cohort, we found that *HOXB13* gene expression was positively correlated with HER2 protein expression in estrogen receptor-positive (ER+) but not ER-negative (ER-) tumors^8^. Considering the relative paucity of data on *HOXB13* expression in HER2+ and ER- breast cancer, we sought to examine *HOXB13* expression patterns in independent cohorts. ^8,18–23^

## Methods

### Data and analysis

#### Transcriptomics

Messenger RNA (mRNA) sequencing (RNAseq) data from the TCGA PANCAN atlas breast cancer dataset (https://gdc.cancer.gov/about-data/publications/pancanatlas/EBPlusPlusAdjustPANCAN_IlluminaHiSeq_RNASeqV2.geneExp.tsv) were used to correlate *HOXB13* gene expression with immunohistochemical (IHC)-based HER2 classification in ER+ (ER+PR+, ER+PR-) and ER- (ER-PR+ and ER-PR-) breast cancers. ER, PR and HER2 immunohistochemistry data were obtained from TCGA pancancer atlas file clinical_PANCAN_patient_with_followup.csv.

#### Copy number

DNA copy number information was obtained from the TCGA pancancer atlas^24^. Absolute copy number calls (TCGA_mastercalls.abs_segtabs.txt) and GISTIC thresholded per-gene calls (all_thresholded.by_genes_whitelisted.tsv) were obtained from https://gdc.cancer.gov/about-data/publications/pancanatlas. GISTIC calls were used to identify *ERBB2* and *HOXB13* gene *gains* (indicated by a value of 1) and *amplifications* (indicated by a value of 2) and to separate patients into subgroups. “Gap” tumors were identified as those with *ERBB2* and *HOXB13* GISTIC copy number levels equaling 1 or 2 and *BRCA1* copy number level of 0 or -1. For the validation cohort from Rheinbay et al^25^, the same thresholds for GISTIC calls were used to determine gain and amplification for each gene using the file all_thresholded.by_genes.txt and to separate patients into subgroups. Validation copy number and survival data from the METABRIC breast cancer cohort^26^ and the MSK-IMPACT study were obtained from the cBioPortal website on (https://www.cbioportal.org/study/summary?id=brca_metabric and https://www.cbioportal.org/study/summary?id=msk_impact_2017). Log_2_ tumor/normal copy ratios and cell line information from the Cancer Cell Line Encyclopedia (CCLE) dataset was downloaded from the DepMap portal (version 22Q2, downloaded in October 2023)^27^.

#### Mutational signatures

Mutational signature analysis: Single-nucleotide variant (SNV) mutational signature assignments for TCGA breast cancers were obtained from Polak et al^28^. Signatures were available for a large subset of samples studied here (gap: 43/47 samples; co-gain: 178/191; HER2-only: 78/86). Extracted signatures from Polak et al were associated with known signature names as follows (as provided by the authors): H1: MSI (microsatellite instability); H2: homologous recombination repair deficiency/BRCAness; H3: Aging (cytosine deamination); H4: APOBEC.

The SigProfiler algorithm was used on the copy number (CNV) data from TCGA participants obtained from GDC (https://gdc.cancer.gov/about-data/publications/pancanatlas) to generate mutational signatures (SigProfilerAssignment version 0.1.0, SigProfilerExtractor version 1.1.23, SigProfilerMatrixGenerator version 1.2.19, SigProfilerPlotting version 1.3.18). For the CNV analysis, the matrix tool generateCNVMatrix was used to convert the ABSOLUTE calls supplied by TCGA (TCGA_mastercalls.abs_segtabs.fixed.txt) into the sigProfiler format. sigProfilerAssigment was then used to assign the number of contributing events from each of the 25 CNV signatures to each tumor individually.

*TP53* and *PIK3CA* mutation analysis considered all mutation types as impactful, except those classified as “Silent”, “Intron”, “5’UTR”, “3’UTR”, and “IGR” (intergenic region).

#### Fluorescence In Situ Hybridization

For validation of findings, formalin-fixed paraffin-embedded (FFPE) tumor samples from 79 consecutive *HER2*-amplified breast cancer patients diagnosed between 2010-2013 at Massachusetts General Hospital were retrospectively collected under IRB protocol 2002-P002059/MGH. Clinical determination of hormone receptor status and *HER2-*amplification status in the 79-patient validation cohort was performed at the Clinical Laboratory Improvement Amendments (CLIA)-certified MGH Clinical Immunohistochemistry and MGH Center for Integrated Diagnostics laboratories following standard protocols using monoclonal antibody clone 6F11 for ER and clone 16 for PR (Leica Microsystems, Inc. Buffalo Grove, IL, USA), and PathVysion *HER-2* DNA probe kit (Abbott, Abbott Park, IL USA). Fluorescence in situ hybridization (FISH) using formalin-fixed paraffin-embedded (FFPE) tumor specimens was employed to analyze *HER2* and *HOXB13* gene amplification status. Briefly, 5-micron sections of FFPE tumor material were prepared, and an H&E section reviewed to select regions for hybridization that contain most tumor cells. A tri-color FISH assay was performed using the using a probe specific to the chromosome 17q *HER2* locus, a probe specific to the *HOXB13* locus and a copy number control probe recognizing centromere 17 (chromosome enumeration probe 17, CEP17). Signal quantitation was used to generate *HER2*/centromere 17 and *HOXB13*/centromere 17 ratios. A ratio of > 2.0 *HER2* and *HOXB13* to CEP17 signals in at least 60 interphase tumor cell nuclei was considered as amplification of *HER2* and *HOXB13*.

#### Survival analysis

TCGA pancancer atlas progression-free survival (PFS) (TCGA-CDR-SupplementalTableS1.xlsx^29^) was used for survival analyses. Kaplan-Meier survival estimates and statistics were calculated with the Python Lifelines package (Version 0.27.8).

#### Statistics

Comparisons between two distributions were performed using the non-parametric Mann-Whitney U test. Multiple-group comparisons were made with the non-parametric Kruskal-Wallis test and Fisher’s exact test for contingency tables. Chi squared tests were used for categorical tests.

##### Visualizations

All graphs were generated from custom Python scripts using the Seaborn package^30^ and the Matplotlib package^31^ Complex structural somatic variations were visualized with Circos^32^ and gTrack (https://github.com/mskilab-org/gTrack version 0.1.0).

##### Code availability

Custom analysis scripts are available under https://github.com/rheinbaylab/Mitsiades_HOXB13_2024.

## Results

### HOXB13 and HER2 gene expression in HER2 positive breast cancer

We have previously demonstrated that *HOXB13* mRNA expression is positively correlated with HER2 immunohistochemistry (IHC) positivity in ER+ but not ER- breast cancers^8^. To further expand upon these findings, we investigated the correlation of *HOXB13* gene expression with IHC-based HER2 protein expression in the TCGA breast cancer cohort. Paired gene expression (mRNA-seq), ER and HER2 IHC-based protein expression and outcome data were available for 707 TCGA breast cancer samples^33^. *HOXB13* gene expression was correlated with HER2 IHC status with significantly higher *HOXB13* mRNA levels in HER+ vs. HER2- breast cancers, suggesting a potential link between these two genes (**Figure 1A**; P=7.98 x 10^-7^). Among HER2+ tumors, *HOXB13* expression was significantly higher in the ER- vs. ER+ subset (**Figure 1B**; P=7.29 x 10^-3^), and among HER2- tumors *HOXB13* expression was also higher in ER- vs ER+ subset (**Figure 1C**; P=7.48 x 10^-5^), suggesting generally higher expression of *HOXB13* in ER- disease.

**Figure 1.**
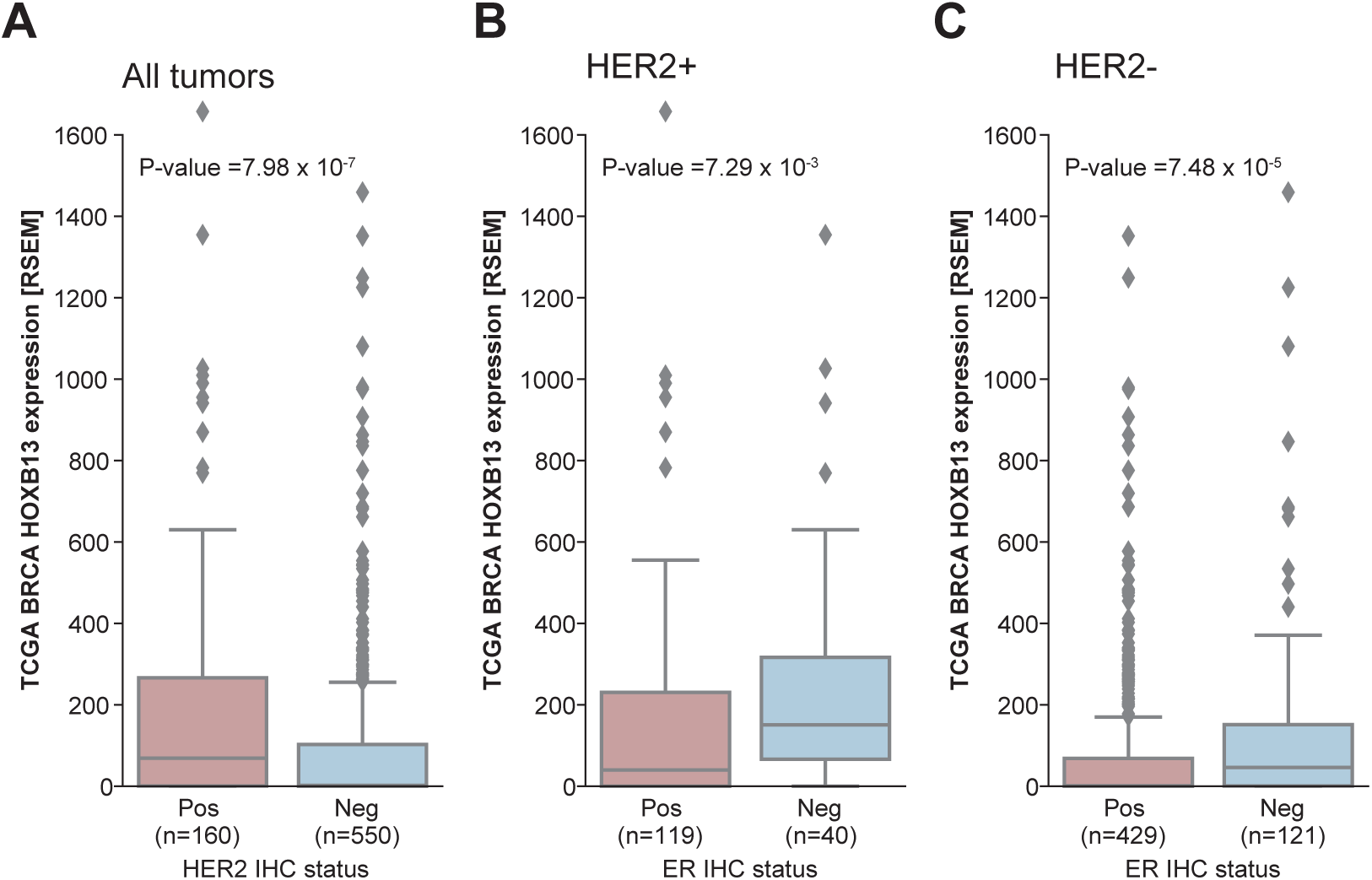
*HOXB13* expression is increased in HER2-positive breast cancer. **(A)***HOXB13* messenger RNA (mRNA) sequencing (RNA-seq) data from the TCGA breast cancer dataset in tumors positive or negative for HER2 by immunohistochemical (IHC). *P*-values calculate with the Wilcoxon two-sample test. Direct comparison of *HOXB13* mRNA in ER+ and ER- HER2+ **(B)** and HER2- **(C)** breast tumors.

### Joint genomic gains of the ERBB2 locus and HOXB13 in a subset of breast cancers

The majority of IHC HER2+ breast tumors is driven by the somatic acquisition of additional copies of the *ERBB2* (encoding the HER2 protein) locus on chromosome 17, either through broad, low-copy arm-level gains or high-level focal amplification, including those caused by chromothripsis^34–36^. The *HOXB13* gene is located 9 Mb downstream of the *ERBB2* locus on chromosome 17q. Thus, the relatively close proximity of these two genes suggests that increased *HOXB13* expression in HER2+ tumors might be due to simultaneous *ERBB2/HOXB13* gene gain. To test this hypothesis, we interrogated absolute copy number calls from the TCGA breast cancer cohort for *HOXB13* and *ERBB2*. We first confirmed that IHC HER2+ breast cancers were enriched for additional *ERBB2* gene copies. Among IHC HER2+ tumors (133), 19% (n=25) had low-level (3-4 total copies) *ERBB2* gains and 52% (n = 69) had high-level amplifications (≥ 5 copies) (**Figure 2A**; interestingly, 31% of IHC HER2- cases also had >2 *ERBB2* copies). Consistent with our hypothesis, *HOXB13* low-level gains or high-level amplifications were enriched in IHC HER2+ cases: 53% of IHC HER2+ tumors had *HOXB13* gains or amplifications, compared to 38% of IHC HER2- tumors **(Figure 2B**; Fisher’s Exact *P* = 8.41 x 10^-4^). For further analyses, we used the TCGA GISTIC-derived copy number calls, as they incorporate overall tumor ploidy in the thresholded gene-level copy number assessment^37^. Following the GISTIC definitions, we classified *ERRB2* and *HOXB13* copy number calls with 1 as “gained”, 2 as “amplified”, and 0 as “unaltered”. Independent of IHC HER2 status, *ERBB2* and *HOXB13* genes were frequently concurrently gained or amplified in the TCGA breast cancer cohort (**Figure 2C**; Fisher’s Exact *P*=2 x 10^-94^). Together, *HOXB13* and *ERBB2* were jointly gained at low level or co-amplified at high-level in 240 participants (22.6%; **Figure 2C**). *ERBB2* without *HOXB13* was gained or amplified in 87 cases (8.2%, **Figure 2C**), and *HOXB13* without *ERBB2* was gained or amplified in 67 tumors (6.3%; **Figure 2C)**. Similarly to TCGA, *HOXB13* gains and amplifications were enriched in HER2+ tumors from METABRIC (**Supplementary Figure 1A-C;** Fisher’s *P* = 0.0018^26^), supporting that *ERBB2/HOXB13* joint gains and amplifications are common in breast cancer. To further validate joint *ERBB2/HOXB13* copy number changes, we performed tri-color fluorescence *in situ* hybridization (FISH) for *ERBB2*, *HOXB13*, and *CEP17* in an independent retrospective consecutive cases series of 79 HER2 IHC3+, HER2-amplified (*ERBB2/CEP17* genomic ratio ≥ 2) breast cancers diagnosed at MGH (**Figure 2D,E)**. Consistent with the TCGA and METABRIC data, *HOXB13* was concurrently gained or co-amplified in 18 of 79 (23%) of *ERBB2*-amplified cases (**Figure 2F**). In the institutional cohort, joint copy gains/amplifications were more frequent in ER+ (24%) than ER- tumors (18%), although this difference was not significant (Fisher’s Exact *P* = 0.5) but mirrored the percentages in the TCGA cohort (24% in ER+, 16% ER-, *P* = 0.02), and trend in the METABRIC cohort (11% in ER+, 8% ER-, p=0.04). In summary, these data demonstrate that *HOXB13* is frequently concurrently gained or amplified with the *ERBB2* gene locus in breast cancer.

**Figure 2.**
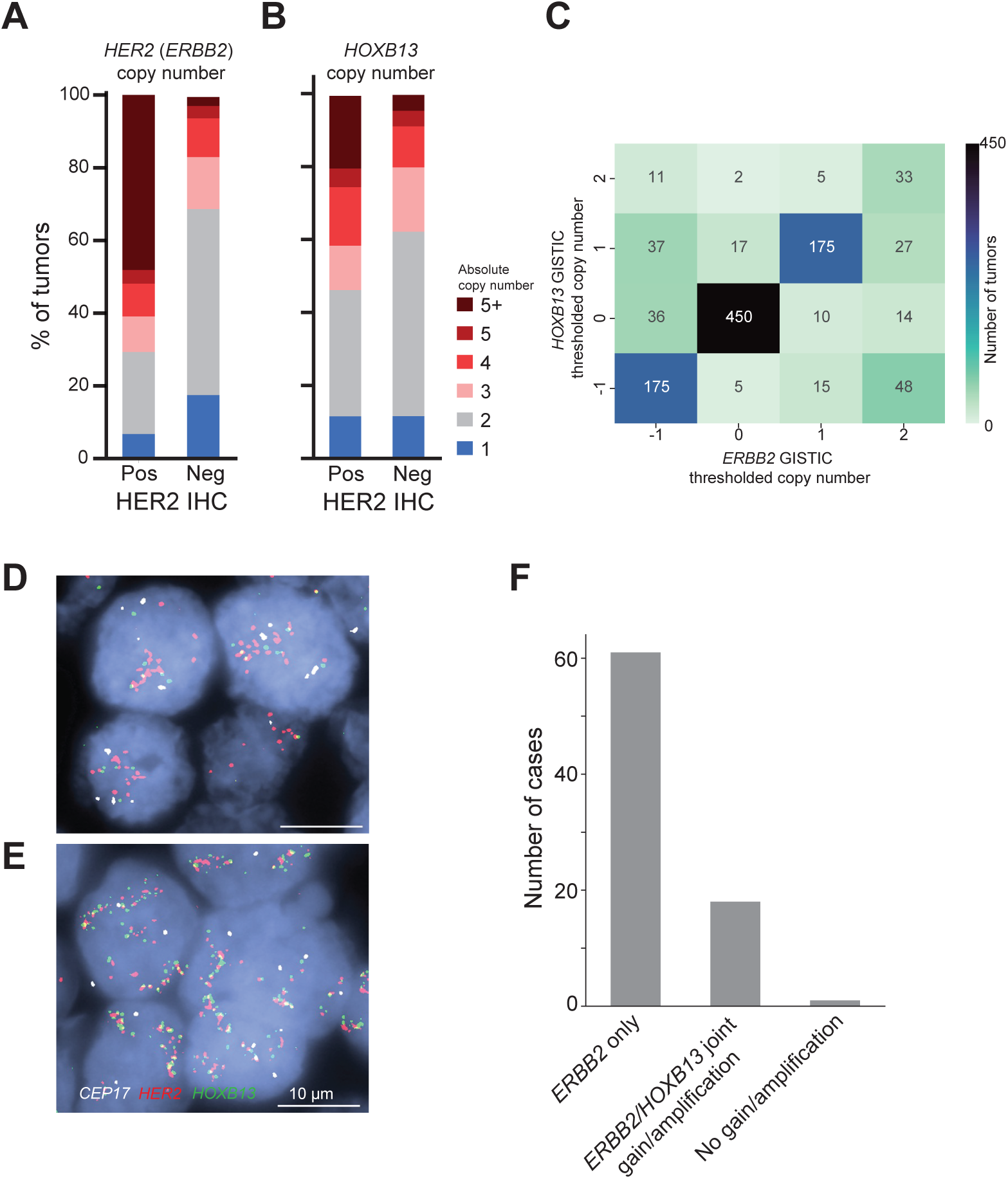
*ERBB2* and *HOXB13* copy gains and amplification in breast cancer. **(A)**Absolute *ERRB2* copy number for IHC HER2+ and HER- negative tumors. **(B)** Absolute *HOXB13* copy number for IHC HER2+ and HER- negative tumors. **(C)** Heatmap of *ERRB2* copy status (x-axis) vs. *HOXB13* copy status (y-axis). Each cell contains the number of tumors with a given *ERRB2/HOXB13* copy number combination. Color intensity scaled with the number of tumors in each cell. **(D)** Photomicrography of representative examples of tri-color FISH assay demonstrating *ERBB2* loci (red), *HOXB13 loci* (green) and centromere 17 control loci (white). Breast cancer showing amplification of *ERBB2 (HER2)* gene only and polysomy of *HOXB13* with an *ERBB2* to *Cep17* ratio > 2, while *HOXB13* to *Cep17* ratio of less than 2. **(E)** Breast cancer samples showing both *ERBB2* and *HOXB13* amplification with ratios of *Cep17* >2 for both genes. Scale bar, right low corner, 10 µm. **(F)** *ERBB2/CEP17* copy ratio for all cases from the institutional cohort in **(D and E).** *P*-value calculated with the Mann-Whitney U test.

### High-level ERBB2/HOXB13 co-amplification is associated with BRCA1 loss

We next investigated the structure of copy gains and amplifications involving the *ERBB2* and *HOXB13* loci. As low-level copy gains typically represent broader, sometimes chromosome-arm sized events, and high-level amplifications tend to be of shorter, focal size, we compared the length distributions of genomic segments for *ERBB2* and *HOXB13* in cases with co-gains and amplifications. Consistent with prior literature, we found that low-level gains generally involved long genomic segments (median length 44 Mb anchored on *ERBB2* and 36 Mb anchored on *HOXB13*), while segments amplified with high copy number were much shorter (median length 0.95 Mb for *ERBB2* and 1.4 Mb for *HOXB13*; **Figure 3A**). The length distribution of gained segments, with a median much larger than the distance (9 Mb) between the two genes, suggests that *ERBB2* and *HOXB13* are jointly amplified through gains of much or all of the chromosome 17q arm. In contrast, the distribution of amplified segments with sizes much less than the 9 Mb genomic distance between the two gene loci suggests a non-contiguous, complex pattern for *ERBB2/HOXB13* high-level co-amplification. Supporting this hypothesis, a complex alternating pattern of amplification of the *ERBB2* and *HOXB13* with a relative interstitial loss (“gap”) between the two loci was apparent in linear genome space in a subset of tumors with *ERBB2/HOXB13* co-amplification (**Figure 3B**). Sometimes this complex event also involved additional chromosomes (e.g. **Figure 3C,D; REFs** ^35,38,39^). This complex event bringing the *ERBB2* and *HOXB13* loci in close proximity is also seen in a representative FISH image, showing juxtaposition (yellow) of the *ERBB2* (red) and *HOXB13* (green) probes (**Figure 3E**). In TCGA breast tumors, the gap created between the *ERBB2* and *HOXB13* amplifications ranged in size from 1.4 Mb to 9 Mb and encompassed up to 423 genes, including many keratin genes, the transcription factors *STAT3*, *STAT5A* and *STAT5B*, and the tumor suppressor *BRCA1.* In contrast, representative tumors with *ERBB2* but no *HOXB13* amplification had comparatively simple chromosomal structure around the *ERBB2* locus and no *BRCA1* loss (**Figure 3F**). To more deeply understand the consequences of this complex rearrangement of *ERRB2* and *HOXB13*, we stratified TCGA tumors by selecting those for further analysis if they had: 1.) gain or amplification of *ERRB2* and *HOXB13* in the presence of relative interstitial loss (defined as -1 or 0 by GISTIC; Methods) of *BRCA1* (“gap”; 48 tumors) 2.) gain/amplification of *ERBB2* and *HOXB13* with no loss of *BRCA1* (“co-gained”; 182 tumors), and 3.) gain/amplification of *ERBB2* and no concurrent gain or amplification of *HOXB13* or *BRCA1* (“focal *ERBB2* amp”; 87 tumors; **Figure 3G; Supplementary Table 1**).

**Figure 3.**
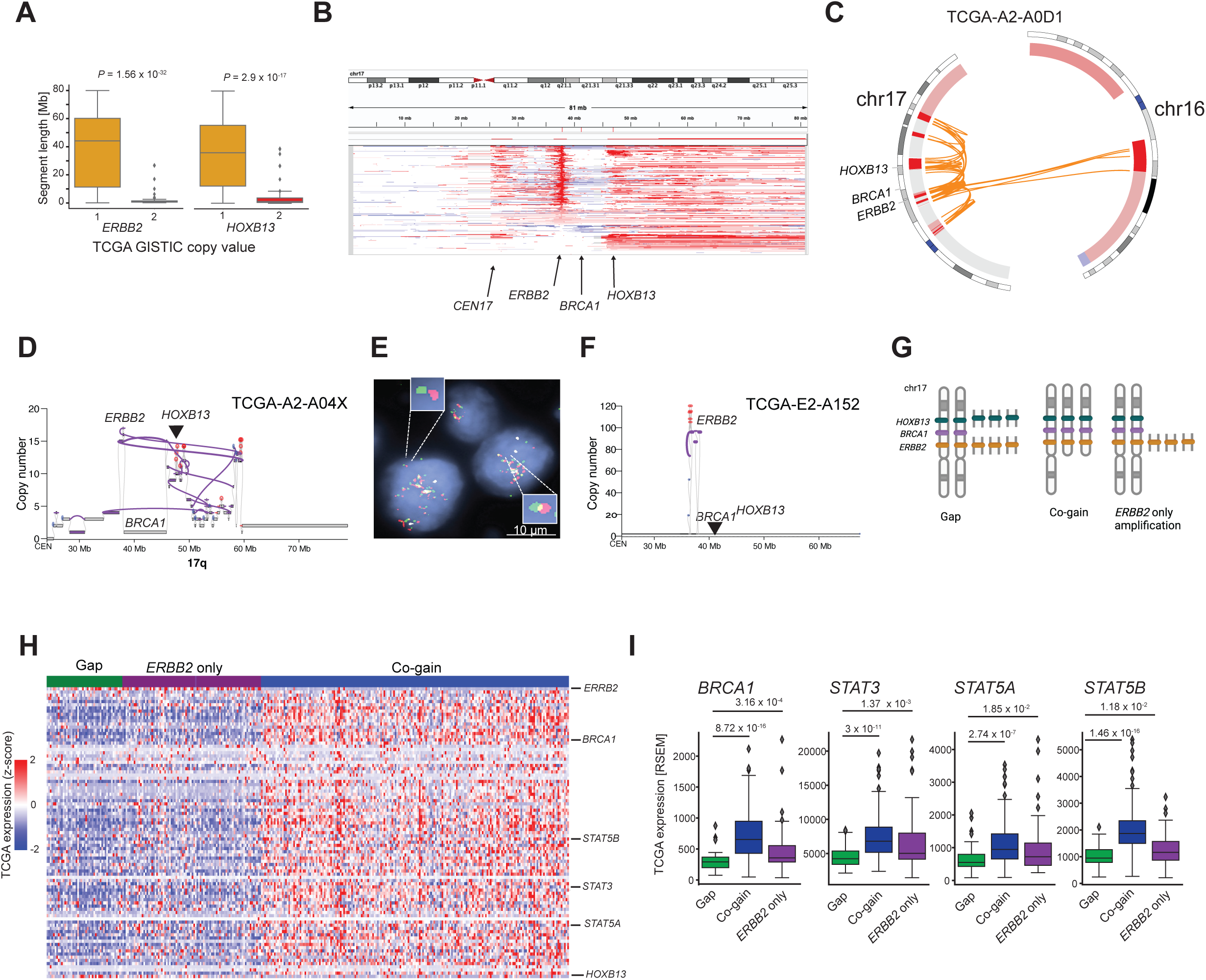
Complex *ERBB2/HOXB13* rearrangements in breast cancer. **(A)**Genomic segment length distribution for *ERBB2* (left) and *HOXB13* (right) for low (TCGA copy status 1) and high (copy status 2) level copy gains. P-values calculated with the Mann-Whitney U test. **(B)** IGV genome viewer^54^ screenshot of chromosome 17 depicting copy number alterations in representative sampling of TCGA breast tumors. White: neutral. Red: copy gain. Blue: copy loss. **(C)** Circos^32^ plot showing complex genomic rearrangements between chromosomal loci encompassing *ERBB2*, *HOXB13*, and *BRCA1* (orange lines) and relative copy number for TCGA tumor TCGA-A2-A0D1 (ER-/PR-/HER2+). **(D)** Example of a complex rearrangement including *ERRB2*, *HOXB13* and loss of *BRCA1*. **(E)** Representative high-power image of FISH assay from a tumor cell demonstrating spatially distinct *ERBB2* (red probe) and *HOXB13* (green probe) loci (enlarged insert image, upper left), and spatially overlapping *ERBB2* and *HOXB13* loci (yellow) consistent with interstitial deletion (enlarged insert image, bottom right). **(F)** Example of structural variants and copy number events in a focal *ERBB2* tumor (no *HOXB13* gain or amplification). **(G)** Cartoon illustrating copy number states of co-gain (left), focal *ERBB2* (middle) and gap (right) tumors. Cartoon created with Biorender.com. **(H)** Gene expression heatmap of *ERRB2* (top), *HOXB13* (bottom), and genes located between these genes in linear genome space. Values are row-normalized for each gene. (**I**) Gene expression values for select genes from **(H)** by copy number category. Individual points denote tumor samples. Boxes indicate median and interquartile range. *P*-values calculated with the Mann-Whitney U test.

Consistent with the observed genomic structure, mRNA expression of genes located inside the “gap”, including *STAT3, STAT5A, STAT5B,* and *BRCA1*, was significantly diminished compared to focal *HER2-*only amplified and *ERBB2/HOXB13* co-gained tumors (**Figure 3H,I**). Importantly, *BRCA1* expression was decreased to an average level similar to triple-negative TCGA breast tumors, many of which are driven by a *BRCA1*-loss phenotype^28^ (**Supplementary Figure 2A**). Notably, we did not observe evidence for additional *BRCA1* somatic mutation or epigenetic silencing events in *ERBB2/HOXB13* co-amplified tumors^28^, suggesting that *BRCA1* could be a secondary driver in *ERBB2/HOXB13* amplified cases through potential haploinsufficiency^38^. Multiple genomic rearrangement mechanisms have been proposed to underlie *ERBB2* amplification in breast cancer^35,40–42^. Our findings suggest that there are at least two distinct classes of high-level *ERBB2* amplification events in breast cancer, *ERBB2/HOXB13* high-level co-amplification with relative loss of genes located between them (including *STAT3/5* and *BRCA1*), and *ERBB2*-only amplification. Importantly, the “gap” rearrangement does not appear to be a specific consequence of loss of genome integrity induced by the frequent mutations observed in the tumor suppressor gene *TP53* in breast cancer: *TP53* loss-of-function mutations were significantly enriched in tumors with focal *ERBB2* mutations compared to gap or co-gain tumors (Fisher’s Exact P=0.074 for focal *ERBB2* only vs gap and Fisher’s Exact P=1.52 x 10^-7^ for *ERBB2* vs co-gain; **Supplementary Figure 2B).**

#### ERBB2/HOXB13 co-amplification appears unique to breast cancer

Because recurrent, focal, *ERBB2* amplification is a known driver in other tumor types, including colon cancer (TCGA code COAD), uterine corpus endometrial carcinoma (UCEC), lung adenocarcinoma (LUAD) and stomach adenocarcinoma (STAD)^43^, we tested whether *ERBB2/HOXB13* complex amplification with relative “gap” was present in these tumor types by comparing the copy ratio of *ERBB2* and *HOXB13* to *BRCA1*. Among *ERBB2*-amplified carcinomas, *ERBB2/HOXB13* co-amplification with relative *BRCA1* loss was not observed in colon, stomach, lung and uterine carcinomas (**Figure 4A**), suggesting that this complex genomic rearrangement is unique to breast cancer. This observation was confirmed in the MSK-IMPACT dataset^44^ of metastatic cancers (**Figure 4B**). Furthermore, genomic sequence analysis of the cancer cell line data from The Cancer Cell Line Encyclopedia (CCLE)^27^ also revealed that the complex *ERBB2/HOXB13* amplification with relative *BRCA1* loss is exclusively found in breast cancer cell lines (**Figure 4C**). Cell lines with this alteration included the commonly studied HER-positive models HCC202 (HER2+/ER-; Her2), BT-474 (HER2+/ER+; LumB), ZR-75-30 (HER2+/ER+; LumB), HCC1419 (HER2+/ER+/-; LumB/Her2), HCC2218 (HER2+/ER-; Her2) and EFM-192 (HER2+/ER+; LumB) (**Figure 4C**)^45^.

**Figure 4.**
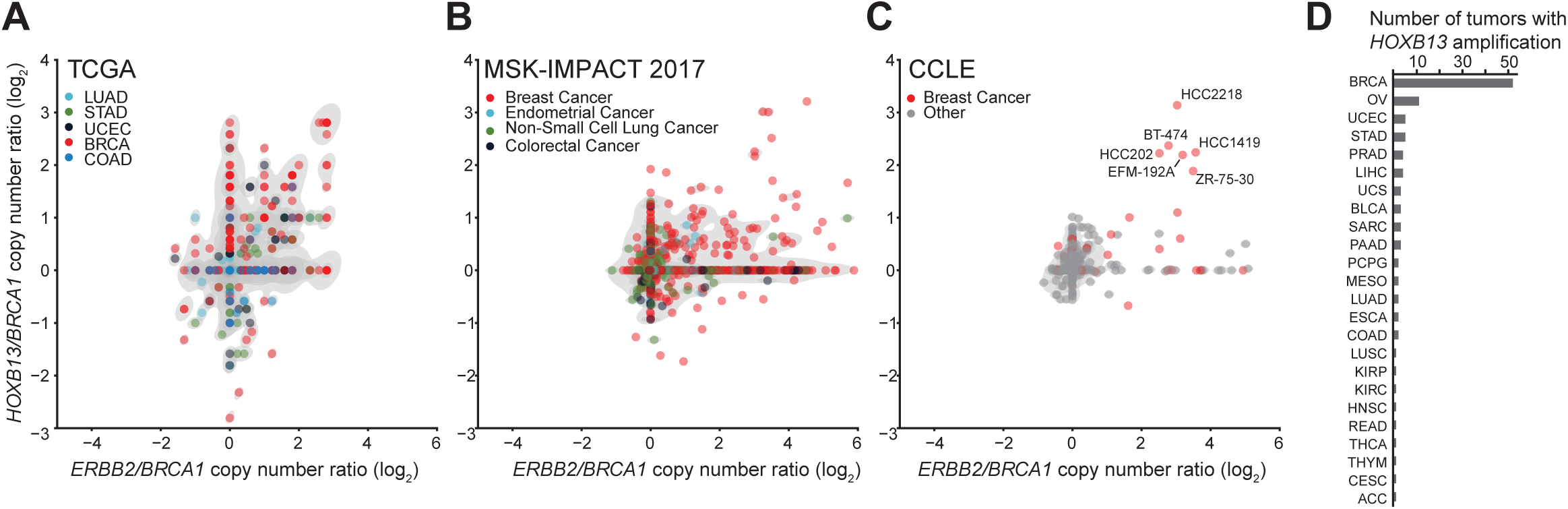
Gap tumors are enriched in breast cancer. **(A)**log2 *HOXB13/BRCA1* vs. *ERBB2/BRCA1* copy number ratio for TCGA cohorts where *ERBB2* amplification is common: UCEC: uterine corpus endometrial carcinoma; STAD: stomach adenocarcinoma; COAD: colon adenocarcinoma; LUAD: lung adenocarcinoma. **(B)** log2 *HOXB13/BRCA1* vs. *ERBB2/BRCA1* copy number ratio for the MSK-IMPACT cohort^44^ and **(C)** for cell lines from the Cancer Cell Line Encyclopedia (CCLE). Breast cancer cell lines are highlighted in red, and gap lines are labeled. **(D)** Number of tumors in the TCGA with *HOXB13* high-level amplification, stratified by cohort. Tumor types indicated by TCGA tumor type code.

Interestingly, high-level amplification of *HOXB13* itself appears restricted to breast cancer in the TCGA cohort (**Figure 4D**), and this alteration is strongly associated with *ERRB2* amplification: 65% of high-level amplifications occur in a background of *ERBB2* amplification (*P*=8.19 x 10^-21^).

Taken together, these results suggest a specific higher-order genomic conformation in breast cells that favors formation of the complex *ERBB2/HOXB13* amplicon, and that the *ERBB2/HOXB13* amplicon with relative *BRCA1* loss confers a specific fitness advantage to breast cancer.

#### Mutational signatures in ERBB2/HOXB13 co-amplified tumors

Enzymes of the APOBEC3 family are active in breast tumors, where they introduce a specific pattern of mutations that can be identified through signature mutational patterns. HER2+ breast cancers specifically have been associated with APOBEC enzyme activity, leaving a characteristic mutational signature in these tumors^46^. Indeed, *ERBB2*-amplified tumors (either focal or with gap) had significantly higher APOBEC mutational signature contributions than co-gain tumors with slightly higher activity in gap over focal *ERBB2*-only genomes (**Figure 5A**). The relative loss of the *BRCA1* tumor suppressor in *ERBB2/HOXB13* co-amplified tumors prompted us to test whether these tumors would show genomic signatures of homologous recombination repair deficiency (HRD)^47–49^. Unexpectedly, we found that the relative contribution of HRD single-nucleotide mutation is significantly lower in “gap” tumors with *BRCA1* loss than either co-gain or *ERBB2*-only amplified tumors (**Figure 5B**). This lower contribution appears to be due at least in part to the higher fractions of mutations attributed to other mutational signatures (including APOBEC), as the total number of HRD-attributed mutations is not significantly different between the three groups (**Figure 5C**). Because HRD can also introduce systematic copy number (CN) changes in cancer cells, especially tandem duplications, we investigated differences in copy number signatures between gap, co-gained and focal *ERBB2* amp tumors^47^. However, we did not observe evidence for differential HRD-associated copy number patterns. Instead, we detected differences in the number of CN5 events between gap and focal *ERBB2* amp tumors (P=0.0004) and in CN5 and CN7 events between gap and co-gain tumors (P = 0.0001 and P=0.0009 respectively (**Figure 5D**). Both CN5 and CN7 are associated with chromothripsis, circular DNA amplicons and poor prognosis^47^. Compared to co-gain tumors, gap and focal *ERBB2* amp tumors had higher copy number of *ERBB2*, which may contribute to the significant differences in the CN7 signature between them. Gap tumors had significantly more chromothripsis-associated signature CN5 than the focal *ERBB2* amp tumors, suggesting that different mechanisms underlie the structures of these amplicons. Interestingly, CN7 is also enriched in tumors from Black and Asian participants^47^. We therefore tested whether the *ERBB2/HOXB13* co-amplified gap structure was similarly enriched in participants from TCGA. Compared to focal *ERBB2* amp tumors or *ERBB2/HOXB13* co-gained tumors, tumors with gap were significantly more common in participants that identified as Asian (**Figure 5E; P=0.0008**), a finding we confirmed in a separate cohort of breast tumors from different ancestries^25^ (**Supplementary Figure 3**). This finding is consistent and expands upon prior reports that HER2+ tumors are enriched in this population^50,51^.

**Figure 5.**
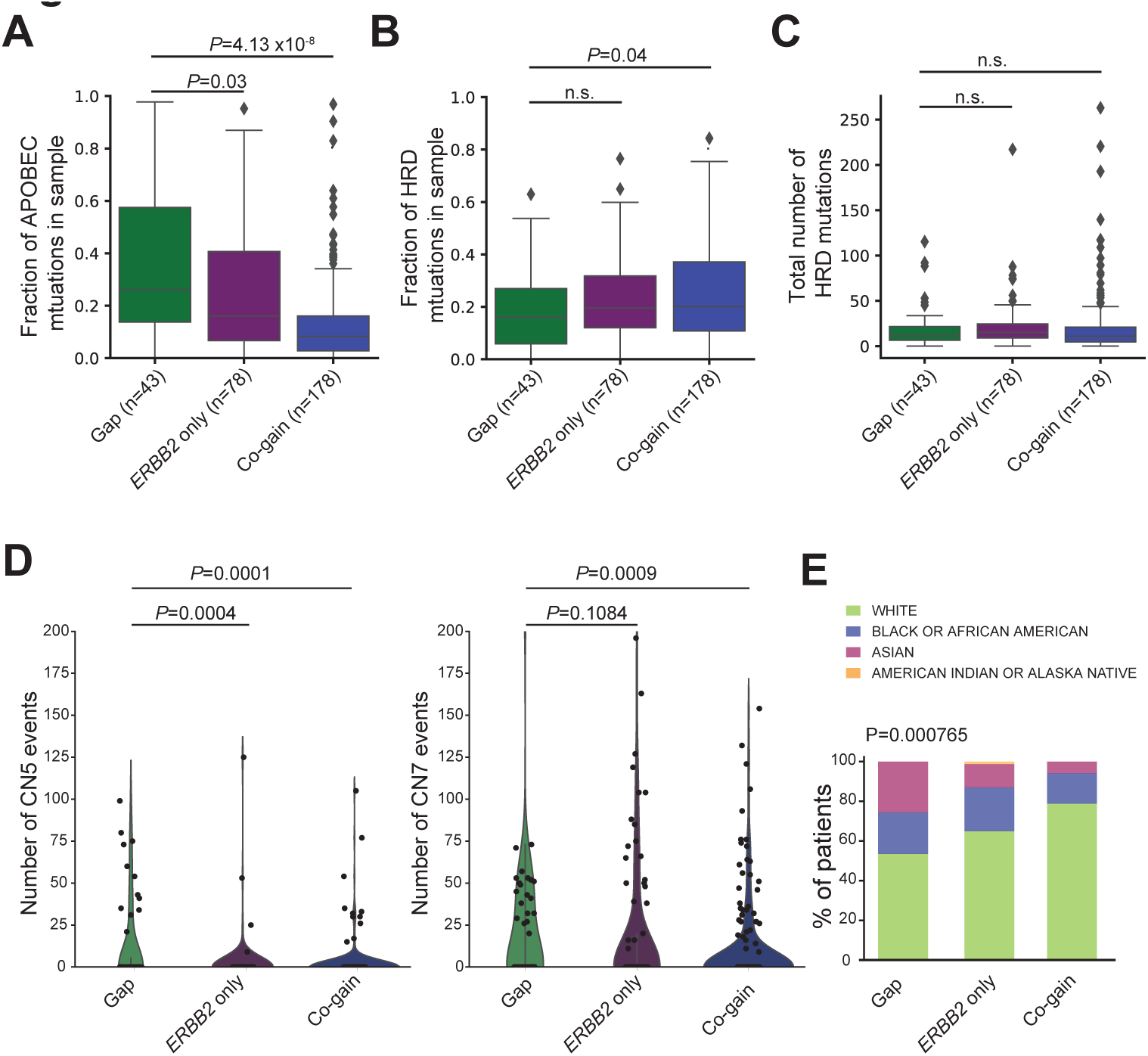
**Mutational signature analysis of *ERBB2/HOXB13* amplified tumors.** Fraction of total mutations attributed to the **(A)** APOBEC or **(B)** homologous recombination repair (HRD) mutational signature. Each dot denotes a tumor sample. Boxplots indicate median and interquartile range. P-values calculated with the non-parametric Mann-Whitney U test. **(C)** Number of total mutations attributed to the HRD signature in tumors from each category. **(D)** Distribution of copy number events attributed to copy number signatures CN5 (left) and CN7 (right). P-values calculated with the Mann-Whitney U test. **(E)** *ERBB2/HOXB13* copy number gains by self-reported race from TCGA. Gap tumors are significantly more common in Asian participants.

It is important to note that the relative dearth of data from exome sequencing limits robust detection of HRD from genomic scars, copy number patterns and mutational signatures. Therefore, we cannot exclude the possibility of differential HRD between the “gap” and ERRB2-only or co-gain tumors. Yet the lack of strong HRD-related point mutation patterns suggests that *ERBB2/HOXB13* tumors are driven by other oncogenes, including amplified *ERBB2* and *HOXB13*, and potentially others.

#### Differential outcome between patients with ERBB2/HER13 and focal ERBB2 amplified tumors

We next examined whether there are outcome differences between *ERBB2/HOXB13* co-gained, *ERBB2/HOXB13*-gap and focal *ERBB2* amplified cases. To increase statistical power due to small numbers, we combined patients from the TCGA and METABRIC cohorts. Among all patients, progression-free survival (PFS) was substantially different between patients with *ERBB2/HOXB13* gap tumors, focal *ERBB2* only amplifications and low-level *ERBB2/HOXB13* co-gains, with gap tumors associated with inferior PFS (HR = 3.3; *P* = 0.068 between gap and focal *ERBB2*-only and HR = 5.2, P=0.022 between gap and co-gain; **Figure 6A**). There was no significant difference in PFS (*P* = 0.66) in patients with focal *ERBB2*-only tumors and those with *ERBB2/HOXB13* co-gain tumors (**Figure 6A**), and there was no difference between the three *ERBB2/HOXB13* groups in ER+ tumors. However, in ER- tumors, we found significantly worse PFS for patients with gap compared to focal *ERBB2*-only tumors (*P* = 0.0078), with median PFS of 999 days for patients with gap tumors compared to 6058 days for those with *ERBB2*-only amplified tumors (**Figure 6B**). No significant difference in PFS was observed between the ER+ or ER- *ERBB2/HOXB13* co-gained and gap or focal *ERBB2* amplified tumors.

**Figure 6.**
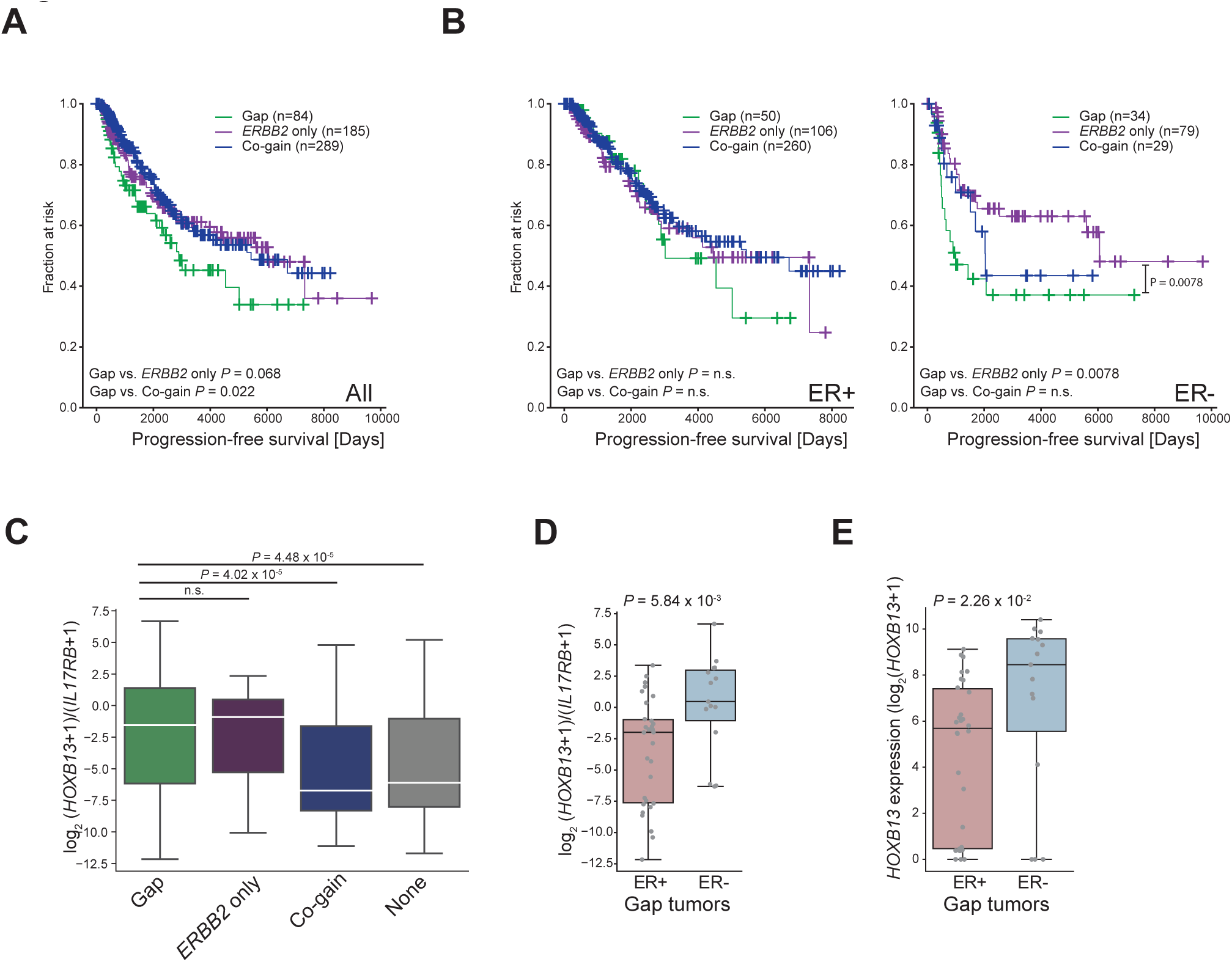
Association of *ERRB2* and *HOXB13* copy number with outcome. **(A)**Progression-free survival of participants from the TCGA BRCA cohort, stratified by tumor copy number category. Censored data are indicated by cross bars. **(B)** Progression-free survival of TCGA participants with IHC ER+ (left) and IHC ER- (right) tumors. **(C)** *HOXB13/IL17RB* gene expression ratio (log_2_+1) by category. P-values calculated with the Mann-Whitney U test. **(D)** *HOXB13/IL17RB* gene expression ratio (log_2_+1) for IHC ER+ and IHC ER- tumors. P-values calculated with the Mann-Whitney U test.

Because copy number is generally correlated with gene expression, we hypothesized that at least in some tumors *HOXB13* amplification underlies high *HOXB13/IL17RB* scores. Indeed, we observed that the *HOXB13/IL17RB* ratio was significantly higher in gap compared to co-gain tumors (*P* = 0.00004) or those with no *ERBB2/HOXB13* amplification (*P* = 0.000045) but not those with focal *ERBB2* amplification only (**Figure 6C**; *P* = 0.68). Within gap tumors, *HOXB13/IL17RB* expression ratio was significantly higher in ER- vs. ER+ tumors (*P* = 5.84 x 10^-3^; **Figure 6D**), explained at least partially by higher *HOXB13* expression levels in ER- gap tumors overall (**Figure 6E**). Together, these findings suggest a possible connection between the predictive value of the *HOXB13/IL17RB* ratio and *ERBB2/HOXB13* amplification, potentially in conjunction with relative loss of genes in the gap between *ERBB2* and *HOXB13* on chromosome 17. Further study of these individual genes will be necessary to elucidate the exact factors underlying the observed benefit.

## Discussion

The expression of *HOXB13*, the primary determinant of the predictive performance of the BCI biomarker, has been extensively studied in ER+ HER2- breast cancer patients from multi-institutional cohorts as well as randomized clinical trial cohorts of both adjuvant and extended adjuvant hormonal therapy. When treated with anti-hormonal agents, patients whose tumors had high *HOXB13/IL17RB* gene expression ratios experienced a significant reduction in the risk of recurrence with an absolute benefit ranging from 9.7% to 16.5%^2,3,6^. However, the clinical relevance of *HOXB13* expression in patients with ER- and HER2+ tumors is currently not understood. Consistent with our previous studies^8,52^, herein we demonstrate that *HOXB13* expression is significantly higher in tumors from HER2+ versus HER- breast cancer patients from the TCGA, and that ER- tumors generally express more *HOXB13* than ER+ tumors.

Complex genomic events involving *ERBB2* and other regions on chromosome 17, including chromothripsis and formation of double minute chromosomes carrying *ERRB2* are well known^26,34,38,41,53^. Our analysis of the TCGA and METABRIC breast cancer cohorts show that *HOXB13* is frequently co-amplified with *ERBB2* as complex high-level rearrangements with relative interstitial *BRCA1*, *STAT3* and *STAT5A/B* loss. Thus, in addition to estradiol-induced regulation of *HOXB13* gene expression^8^, genomic amplification provides another mechanism of *HOXB13* gene expression modulation, especially in ER-/HER2+ tumors. We further validated low- and high-level co-gain/co-amplification findings by FISH analysis of a consecutive clinical case series of HER2+ breast cancer cases, with compatible frequency of co-amplification/co-gain to those observed in the TCGA and METABRIC cohorts. Interestingly, we observed the *ERBB2/HOXB13* co-amplification with *BRCA1* deletion exclusively in breast cancer, but not other tumor types with recurrent *ERBB2* amplifications, suggesting that the unique constellation of the complex *ERBB2*-amplicon with its co-amplified and deleted genes confers a specific selective advantage in breast cancer. Consistent with this hypothesis, TCGA and METABRIC breast cancer patients with ER- *ERBB2*/HOXB13-gap tumors displayed significantly shortened progression-free survival. This observation raises the question whether HER2+ breast cancer could be further stratified into genetic subtypes when evaluating resistance patterns in this cancer type and in clinical trials focused on HER2+ tumors. Most importantly, our results also suggest further investigation into whether the *ERBB2*/*HOXB13* gap subtype should be treated more aggressively.

Although *HOXB13* gene expression is the major determinant in the prognostic performance of the BCI HOXB13/IL17RB biomarker in ER+ HER2- breast cancer patients, our current findings suggest that a putative prognostic role for *HOXB13* gene expression in ER- HER2+ patients is likely more complex. We demonstrated ER- *ERBB2/HOXB13*-gap breast cancer patients had a shortened progression-free survival compared to patients with ER- *ERBB2*-only amplified tumors. Although ER- gap tumors had somewhat higher *HOXB13* gene expression compared to ER- *ERRB2*-only tumors, this difference was not significant (**Supplementary Figure 4**). This suggests that *HOXB13* gene expression alone is not associated with inferior progression-free survival in such patients, and that loss of genes in the interstitial gap likely also contribute to the observed phenotype. Although *ERBB2*/HOXB13 gap tumors harbor *BRCA1* loss (without other concomitant *BRCA1* mutations), we were unable to find significant differences in HRD between the tumor subgroups.

Our data therefore suggest that in contrast to HRD-positive, triple-negative breast cancer, *ERBB2/HOXB13* gap tumors may not be sensitive to PARP inhibitors and related therapies, potentially explaining the shortened disease-free survival in patients with these tumors. Future study of the individual genes within the gap between *ERBB2* and *HOXB13* as well as other genes associated with the *HOXB13* amplicon will be necessary to elucidate the relative contribution of these genes to the observed differences in clinical outcome between ER- *ERBB2/HOXB13*-gap and ER- focal *ERBB2* amplified tumors.

In summary, amongst patients with *ERBB2* amplified tumors we identified a subset of cancers with complex co-amplification of *ERBB2* and *HOXB13* with an interstitial deletion that includes *BRCA1*. This complex genomic alteration is unique to breast tumors, and ER- breast cancer patients whose tumors harbor this genomic complex alteration demonstrated shortened progression-free survival.

## Acknowledgements

Funding was provided in part by the Breast Cancer Research Foundation to D.C.S. (BCRF-20-147, BCRF-21-147, BCRF-22-147). E.R. and I.R.M. are partially supported by the RICBAC foundation. We thank Doga Gülhan for help with signature analysis, Michael Lawrence for helpful discussion and Marcin Imielinski for assistance with structural rearrangement visualization.

---

**Supplementary Figure 1:**
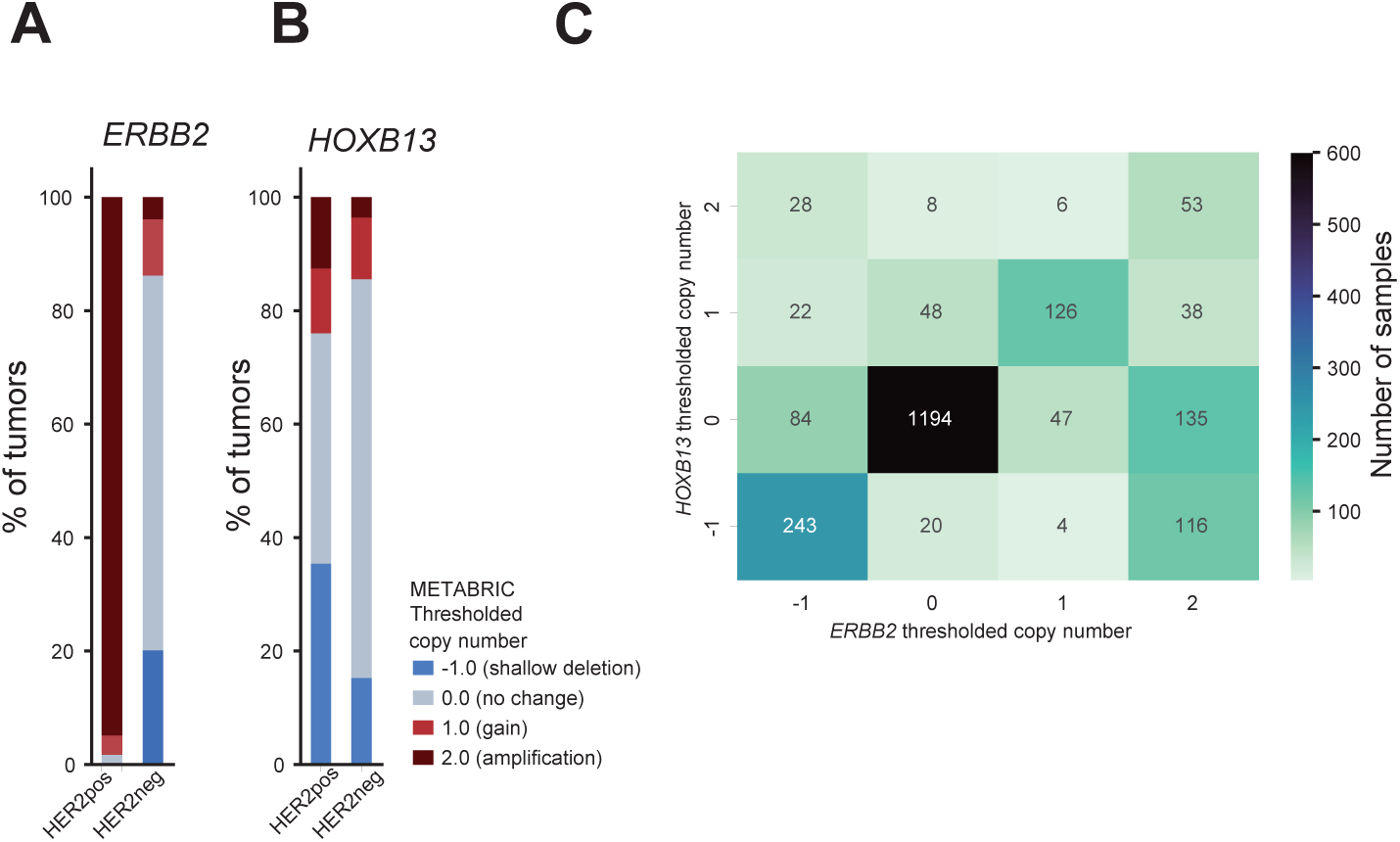
Breakdown of *ERBB2* **(A)** and *HOXB13* **(B)** GISTIC copy number calls in participants from the METABRIC breast cancer cohort^26^, stratified by HER2 status. **(C)** Heatmap of *ERBB2* copy status (x-axis) vs. *HOXB13* copy status (y-axis) for METABRIC samples. Each cell contains the number of tumors with a given *ERBB2/HOXB13* copy number combination. Color intensity scaled with the number of tumors in each cell. **(D)** *ERBB2* absolute copy number from TCGA for *ERBB2/HOXB13* joint gains vs *ERBB2*-only gained/amplified samples.

**Supplementary Figure 2:**
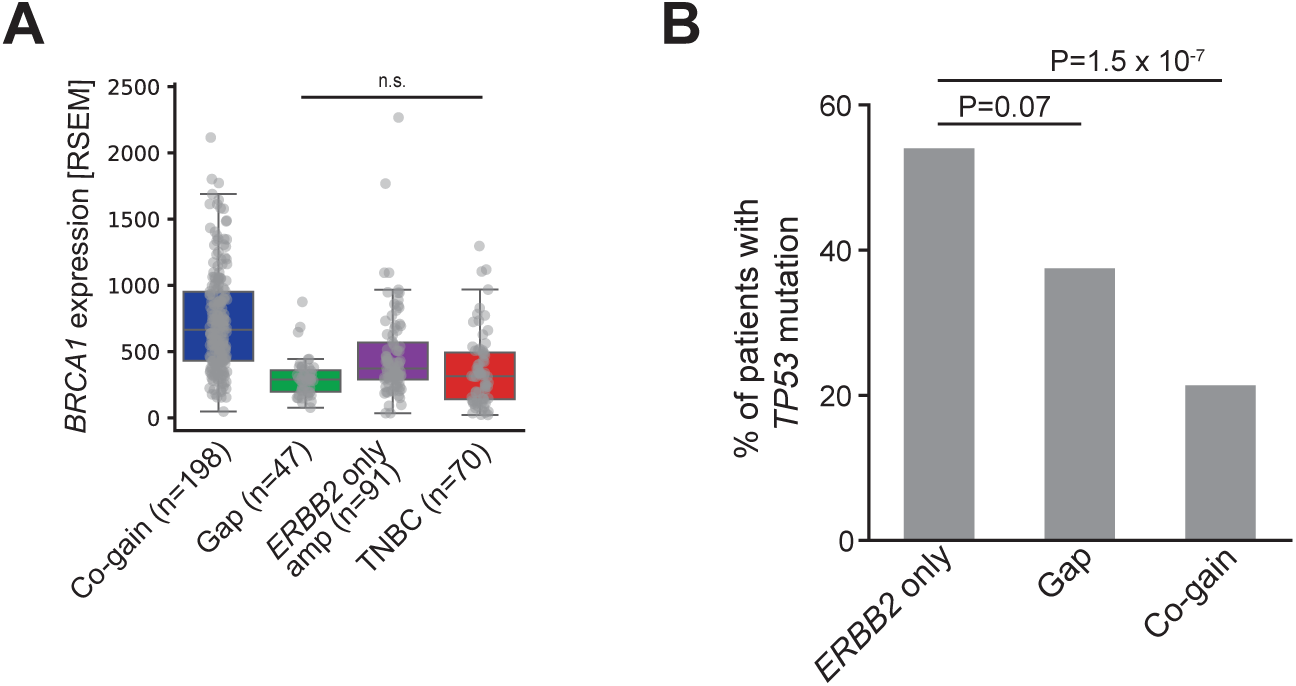
(A) *BRCA1* gene expression in tumors from different *ERBB2/HOXB13* categories. Grey points denote values for individual tumors. Boxplots indicate median and interquartile range (box limits). **(B)** Percentage of tumors with protein-altering *TP53* gene mutations, in co-gain, gap and focal *ERBB2* tumors from the TCGA breast cancer (BRCA) cohort. Pairwise P-values calculated with the Fisher’s exact test.

**Supplementary Figure 3:**
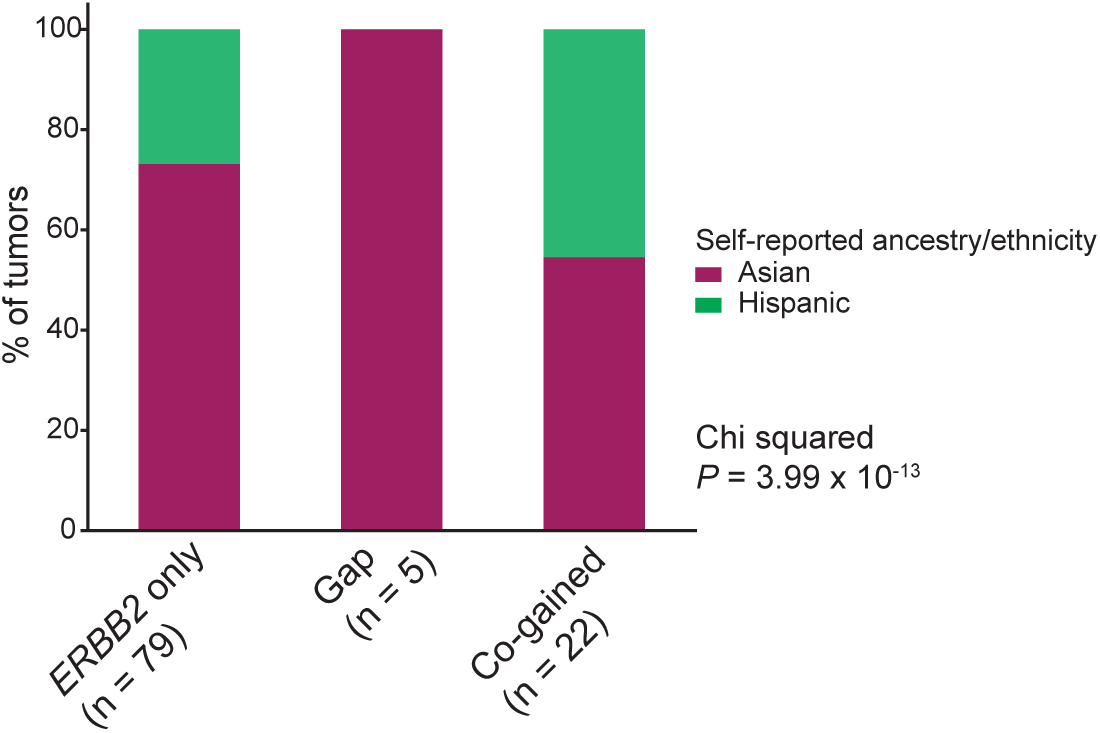
*ERBB2/HOXB13* gain/amplification patterns stratified by ancestry/ethnicity from Rheinbay et al, 2017^25^ confirms enrichment of the complex gap amplicon in tumors from Asian donors.

**Supplementary Figure 4:**
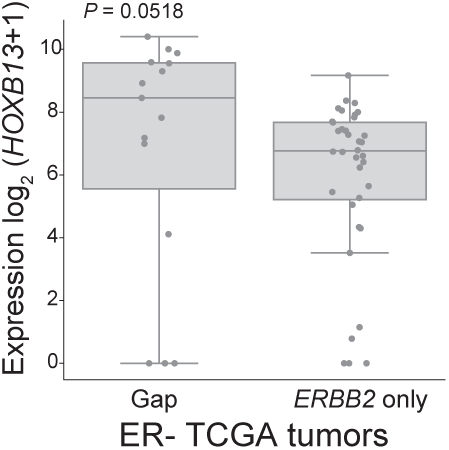
ER- gap tumors have a trend towards higher *HOXB13* mRNA expression compared to ER- *ERBB2* focal amplification tumors (P = 0.0518).

## References

1. Bartlett JMS, Sgroi DC, Treuner K, et al. Breast Cancer Index and prediction of benefit from extended endocrine therapy in breast cancer patients treated in the Adjuvant Tamoxifen-To Offer More? (aTTom) trial. Ann Oncol 2019; 30(11): 1776–83.

2. Bartlett JMS, Sgroi DC, Treuner K, et al. Breast Cancer Index is a predictive biomarker of treatment benefit and outcome from extended tamoxifen therapy: final analysis of the Trans-aTTom study. Clin Cancer Res 2022.

3. Sgroi DC, Carney E, Zarrella E, et al. Prediction of late disease recurrence and extended adjuvant letrozole benefit by the HOXB13/IL17BR biomarker. J Natl Cancer Inst 2013; 105(14): 1036–42.

4. Zhang Y, Schnabel CA, Schroeder BE, et al. Breast cancer index identifies early-stage estrogen receptor-positive breast cancer patients at risk for early- and late-distant recurrence. Clin Cancer Res 2013; 19(15): 4196–205.

5. Sgroi DC, Sestak I, Cuzick J, et al. Prediction of late distant recurrence in patients with oestrogen-receptor-positive breast cancer: a prospective comparison of the breast-cancer index (BCI) assay, 21-gene recurrence score, and IHC4 in the TransATAC study population. Lancet Oncol 2013; 14(11): 1067–76.

6. Noordhoek I, Treuner K, Putter H, et al. Breast Cancer Index Predicts Extended Endocrine Benefit to Individualize Selection of Patients with HR(+) Early-stage Breast Cancer for 10 Years of Endocrine Therapy. Clin Cancer Res 2021; 27(1): 311–9.

7. Treuner KS, D. Schnabel, C., Benner, C., Heinz S. Characterization of HOXB13-induced estrogen receptor reprogramming in breast cancer cells. Cancer Research 2021; 81(4): PS17–32.

8. Wang Z, Dahiya S, Provencher H, et al. The prognostic biomarkers HOXB13, IL17BR, and CHDH are regulated by estrogen in breast cancer. Clin Cancer Res 2007; 13(21): 6327–34.

9. Cerda-Esteban N, Spagnoli FM. Glimpse into Hox and tale regulation of cell differentiation and reprogramming. Dev Dyn 2014; 243(1): 76–87.

10. Huang L, Pu Y, Hepps D, Danielpour D, Prins GS. Posterior Hox gene expression and differential androgen regulation in the developing and adult rat prostate lobes. Endocrinology 2007; 148(3): 1235–45.

11. Abate-Shen C. Deregulated homeobox gene expression in cancer: cause or consequence? Nat Rev Cancer 2002; 2(10): 777–85.

12. Brechka H, Bhanvadia RR, VanOpstall C, Vander Griend DJ. HOXB13 mutations and binding partners in prostate development and cancer: Function, clinical significance, and future directions. Genes Dis 2017; 4(2): 75–87.

13. Hung YC, Ueda M, Terai Y, et al. Homeobox gene expression and mutation in cervical carcinoma cells. Cancer Sci 2003; 94(5): 437–41.

14. Lopez R, Garrido E, Vazquez G, et al. A subgroup of HOX Abd-B gene is differentially expressed in cervical cancer. Int J Gynecol Cancer 2006; 16(3): 1289–96.

15. Yamashita T, Tazawa S, Yawei Z, et al. Suppression of invasive characteristics by antisense introduction of overexpressed HOX genes in ovarian cancer cells. Int J Oncol 2006; 28(4): 931–8.

16. Zhao Y, Yamashita T, Ishikawa M. Regulation of tumor invasion by HOXB13 gene overexpressed in human endometrial cancer. Oncol Rep 2005; 13(4): 721–6.

17. Miao J, Wang Z, Provencher H, et al. HOXB13 promotes ovarian cancer progression. Proc Natl Acad Sci U S A 2007; 104(43): 17093–8.

18. Arpino G, Wiechmann L, Osborne CK, Schiff R. Crosstalk between the estrogen receptor and the HER tyrosine kinase receptor family: molecular mechanism and clinical implications for endocrine therapy resistance. Endocr Rev 2008; 29(2): 217–33.

19. Montemurro F, Di Cosimo S, Arpino G. Human epidermal growth factor receptor 2 (HER2)-positive and hormone receptor-positive breast cancer: new insights into molecular interactions and clinical implications. Ann Oncol 2013; 24(11): 2715–24.

20. Kurokawa H, Lenferink AE, Simpson JF, et al. Inhibition of HER2/neu (erbB-2) and mitogen-activated protein kinases enhances tamoxifen action against HER2-overexpressing, tamoxifen-resistant breast cancer cells. Cancer Res 2000; 60(20): 5887–94.

21. Schiff R, Massarweh SA, Shou J, et al. Advanced concepts in estrogen receptor biology and breast cancer endocrine resistance: implicated role of growth factor signaling and estrogen receptor coregulators. Cancer Chemother Pharmacol 2005; 56 **Suppl 1**: 10–20.

22. Massarweh S, Osborne CK, Jiang S, et al. Mechanisms of tumor regression and resistance to estrogen deprivation and fulvestrant in a model of estrogen receptor-positive, HER-2/neu-positive breast cancer. Cancer Res 2006; 66(16): 8266–73.

23. Montaser RZ, Coley HM. Crosstalk between ERalpha and Receptor Tyrosine Kinase Signalling and Implications for the Development of Anti-Endocrine Resistance. Cancers (Basel) 2018; 10(6).

24. Taylor AM, Shih J, Ha G, et al. Genomic and Functional Approaches to Understanding Cancer Aneuploidy. Cancer cell 2018; 33(4): 676–89 e3.

25. Rheinbay E, Parasuraman P, Grimsby J, et al. Recurrent and functional regulatory mutations in breast cancer. Nature 2017; 547(7661): 55–60.

26. Curtis C, Shah SP, Chin SF, et al. The genomic and transcriptomic architecture of 2,000 breast tumours reveals novel subgroups. Nature 2012; 486(7403): 346–52.

27. Ghandi M, Huang FW, Jané-Valbuena J, et al. Next-generation characterization of the Cancer Cell Line Encyclopedia. Nature 2019; 569(7757): 503–8.

28. Polak P, Kim J, Braunstein LZ, et al. A mutational signature reveals alterations underlying deficient homologous recombination repair in breast cancer. Nature genetics 2017; 49(10): 1476–86.

29. Liu J, Lichtenberg T, Hoadley KA, et al. An Integrated TCGA Pan-Cancer Clinical Data Resource to Drive High-Quality Survival Outcome Analytics. Cell 2018; 173(2): 400–16 e11.

30. Waskom ML. Seaborn: statistical data visualization. Journal of Open Source Software 2021; 6(60): 3021.

31. Hunter JD. Matplotlib: A 2D graphics environment. Computing in science & engineering 2007; 9(03): 90–5.

32. Krzywinski M, Schein J, Birol I, et al. Circos: an information aesthetic for comparative genomics. Genome research 2009; 19(9): 1639–45.

33. Cancer Genome Atlas N. Comprehensive molecular portraits of human breast tumours. Nature 2012; 490(7418): 61–70.

34. Lin CL, Tan X, Chen M, et al. ERalpha-related chromothripsis enhances concordant gene transcription on chromosome 17q11.1-q24.1 in luminal breast cancer. BMC Med Genomics 2020; 13(1): 69.

35. Cortes-Ciriano I, Lee JJ, Xi R, et al. Comprehensive analysis of chromothripsis in 2,658 human cancers using whole-genome sequencing. Nature genetics 2020; 52(3): 331–41.

36. Cai H, Kumar N, Bagheri HC, von Mering C, Robinson MD, Baudis M. Chromothripsis-like patterns are recurring but heterogeneously distributed features in a survey of 22,347 cancer genome screens. BMC genomics 2014; 15: 82.

37. Mermel CH, Schumacher SE, Hill B, Meyerson ML, Beroukhim R, Getz G. GISTIC2.0 facilitates sensitive and confident localization of the targets of focal somatic copy-number alteration in human cancers. Genome Biol 2011; 12(4): R41.

38. Inaki K, Menghi F, Woo XY, et al. Systems consequences of amplicon formation in human breast cancer. Genome research 2014; 24(10): 1559–71.

39. Rankins DL, Jr., Smith GS, Hallford DM. Serum constituents and metabolic hormones in sheep and cattle fed Kochia scoparia hay. J Anim Sci 1991; 69(7): 2941–6.

40. Kim H, Nguyen NP, Turner K, et al. Extrachromosomal DNA is associated with oncogene amplification and poor outcome across multiple cancers. Nature genetics 2020; 52(9): 891–7.

41. Hadi K, Yao X, Behr JM, et al. Distinct Classes of Complex Structural Variation Uncovered across Thousands of Cancer Genome Graphs. Cell 2020; 183(1): 197–210 e32.

42. Przybytkowski E, Lenkiewicz E, Barrett MT, et al. Chromosome-breakage genomic instability and chromothripsis in breast cancer. BMC genomics 2014; 15(1): 579.

43. Mishra R, Hanker AB, Garrett JT. Genomic alterations of ERBB receptors in cancer: clinical implications. Oncotarget 2017; 8(69): 114371–92.

44. Zehir A, Benayed R, Shah RH, et al. Mutational landscape of metastatic cancer revealed from prospective clinical sequencing of 10,000 patients. Nat Med 2017; 23(6): 703–13.

45. Dai X, Cheng H, Bai Z, Li J. Breast Cancer Cell Line Classification and Its Relevance with Breast Tumor Subtyping. J Cancer 2017; 8(16): 3131–41.

46. Roberts SA, Lawrence MS, Klimczak LJ, et al. An APOBEC cytidine deaminase mutagenesis pattern is widespread in human cancers. Nature genetics 2013; 45(9): 970–6.

47. Steele CD, Abbasi A, Islam SMA, et al. Signatures of copy number alterations in human cancer. Nature 2022; 606(7916): 984–91.

48. Alexandrov LB, Kim J, Haradhvala NJ, et al. The repertoire of mutational signatures in human cancer. Nature 2020; 578(7793): 94–101.

49. Alexandrov LB, Nik-Zainal S, Wedge DC, et al. Signatures of mutational processes in human cancer. Nature 2013; 500(7463): 415–21.

50. Pan JW, Zabidi MMA, Ng PS, et al. The molecular landscape of Asian breast cancers reveals clinically relevant population-specific differences. Nature communications 2020; 11(1): 6433.

51. Kan Z, Ding Y, Kim J, et al. Multi-omics profiling of younger Asian breast cancers reveals distinctive molecular signatures. Nature communications 2018; 9(1): 1725.

52. Ma XJ, Hilsenbeck SG, Wang W, et al. The HOXB13:IL17BR expression index is a prognostic factor in early-stage breast cancer. J Clin Oncol 2006; 24(28): 4611–9.

53. Li Y, Roberts ND, Wala JA, et al. Patterns of somatic structural variation in human cancer genomes. Nature 2020; 578(7793): 112–21.

54. Thorvaldsdottir H, Robinson JT, Mesirov JP. Integrative Genomics Viewer (IGV): high-performance genomics data visualization and exploration. Briefings in bioinformatics 2013; 14(2): 178–92.

